# A persistent increase in gut permeability correlates with emotional dysregulation following maternal separation in male and female mice

**DOI:** 10.1101/2025.04.04.647178

**Authors:** Adriana Castro-Zavala, Ana E. Nieto-Nieves, Sonia Melgar-Locatelli, Lidia Medina-Rodríguez, M. Carmen Mañas-Padilla, Estela Castilla-Ortega

## Abstract

Early life stress (ELS) can significantly influence vulnerability to the development of psychiatric disorders in adulthood. One of the most widely used preclinical models for investigating ELS is maternal separation with early weaning (MSEW), which mimics early-life neglect. The objective of this study was to evaluate the impact of ELS induced by MSEW on the emotional behaviour of male and female mice, as well as its relationship with intestinal permeability and neuroinflammatory markers in the hippocampus.

Our results show that MSEW leads to increased anxiety-like behaviours in the adulthood, particularly in females, and exacerbates depression-like behaviours and anhedonia in both sexes. Notably, increased intestinal permeability was observed, which correlated with higher anxiety and depression-like responses, suggesting that gut health plays a crucial role in emotional regulation. These alterations in intestinal permeability were long-lasting, indicating persistent effects on gut function following ELS.

Additionally, we observed that MSEW animals showed higher BDNF expression in the hippocampus, particularly in males. However, we did not find significant differences in the long-term survival of adult-born hippocampal cells, as measured by BrdU+ labeling. Furthermore, upon exposure to MSEW, both sexes showed increased NF-κB protein levels. However, only MSEW male mice exhibited changes in TNF-α and BDNF levels, suggesting a sex-specific regulatory mechanism in response to chronic stress.

The novel contribution of this study is its exploration of intestinal permeability as a mechanism linking ELS to emotional and behavioural dysregulation, particularly anxiety and depression. By showing a long-lasting increase in intestinal permeability and its correlation with mood disorders, our study extends the gut-brain axis hypothesis to ELS. Additionally, the inclusion of both male and female mice offers a more comprehensive understanding of the sex-specific effects of early stress, often overlooked in other studies. These findings suggest that intestinal permeability could serve as a biomarker for stress-related psychiatric conditions.

**HIGHLIGHTS:**

- MSEW increases anxiety and depression-like behaviours, especially in females.
- Maternal separation leads to long-lasting gut permeability changes into adulthood.
- Increased gut permeability directly correlates with anxiety and anhedonia-like responses.
- Neuroinflammation via NF-kB may link gut permeability to mood disorders.
- Differential regulation of early-life stress is observed between males and females.

## INTRODUCTION

Increasing evidence suggests that environmental and biological factors during early life stages, such as early life stress (ELS), can significantly influence vulnerability to the development of psychiatric disorders in adulthood. One of the most widely used preclinical models to investigate ELS is maternal separation with early weaning (MSEW) which mimics early life neglect (George et al., 2010; Baracz et al., 2020; Castro-Zavala et al., 2020, 2021a, 2021b). This model has been extensively used as it replicates conditions of early neglect and has been shown to induce various behavioral and neurobiological alterations. Specifically, this manipulation induces impairments in memory and learning (George et al., 2010; Portero-Tresserra et al., 2018; Planchez et al., 2019), depression-like behaviors such as helplessness (George et al., 2010; Portero-Tresserra et al., 2018; Castro-Zavala et al., 2021b), anxiety (George et al., 2010; Valvassori et al., 2017), high stress reactivity during adulthood (Baracz et al., 2020), anhedonia (Matthews and Robbins, 2003; Gracia-Rubio et al., 2016b; Frank et al., 2019), and increased vulnerability to drug use (Gracia-Rubio et al., 2016a; Portero-Tresserra et al., 2018; Castro-Zavala et al., 2020, 2021a, 2021b). However, the potential mechanisms mediating the long-lasting effects of ELS and MSEW on brain and behaviour are numerous and still under investigation.

In this context, the relationship between ELS and intestinal barrier integrity has gained increasing attention. The gut-brain axis (Cryan et al., 2019) involves bidirectional communication between the gut and the brain, mediated by hormonal, immune, and neural signals, among other factors. This connection suggests that what happens in the gut can directly impact mental health (Mitrea et al., 2022; Varanoske et al., 2022). Research has revealed that alterations in intestinal permeability are related to psychiatric disorders, suggesting that changes in gut microbiota and barrier function may be linked to the development of anxious and depressive behaviours (Stevens et al., 2018; Nikolova et al., 2021; Prospero et al., 2021; Mitrea et al., 2022; Jiang et al., 2023) . This connection highlights the importance of understanding how ELS affects gut health, which in turn can influence mental health.

Specifically, it has been proposed that alterations in the intestinal barrier and gut microbiota could precipitate the onset of anxiety and depression through neuroinflammation, which is characterized by the chronic activation of glial cells such as astrocytes and microglia (Peirce and Alviña, 2019). This neuroinflammation may be explained by changes in the intestine that generate localized inflammation, which, over time, could progress into systemic inflammation (Thevaranjan et al., 2017; Haruwaka et al., 2019), potentially manifesting as neuroinflammation. Moreover, neuroinflammation could be triggered by the passage of endotoxins from the intestine into the bloodstream, which may eventually reach the brain (De Punder and Pruimboom, 2015). Additionally, alterations in intestinal permeability have been linked to changes in the permeability of the blood-brain barrier (BBB), leading to the activation of glial cells (microglia and astrocytes) to protect the BBB (Haruwaka et al., 2019).

Neuroinflammation has been associated with alterations in neuronal function and can negatively affect synaptic plasticity and neurogenesis in the hippocampus, processes essential for adaptation and emotional well-being (DiSabato et al., 2016; Guo et al., 2023). Alterations in neuroinflammatory markers, such as TNF-α and NF-κB, have been observed in various stress conditions and may contribute to hippocampal dysfunction (Koo et al., 2010; Boersma et al., 2011; Imielski et al., 2012; McWhirt et al., 2019).. As mentioned earlier, neuroinflammation also alters synaptic plasticity (Doan et al., 2015; Lima Giacobbo et al., 2019). Alterations in Brain-Derived Neurotrophic Factor (BDNF), a crucial factor synaptic plasticity, learning, and memory, have been linked to stress-related disorders and emotional dysregulation (Doan et al., 2015; Björkholm and Monteggia, 2016),

Given that neuroinflammatory mechanisms and disruptions in gut permeability can influence stress responses, it is crucial to consider potential sex differences in these interactions. Despite advancements in understanding these mechanisms, most studies have focused on the male subjects, leaving a gap in the literature regarding sex differences in the response to early stress and its impact on mental health. It has been observed that hormonal and physiological differences between males and females may influence stress responses and the regulation of affective behaviours (Goel et al., 2014; Leistner and Menke, 2020; Veenit et al., 2021). Recently, a study found differences in stress vulnerability and neuroinflammatory responses between female and male rats, suggesting that sex plays a crucial role in how individuals respond to stressors (San Felipe et al., 2024). Males showed greater vulnerability, evidenced by a delay in motor development and alterations in key proteins involved in neuroinflammation. In contrast, females exhibited a more compensatory response, with an increase in IL-10 levels in the hippocampus, suggesting a protective mechanism against stress-induced neurodegenerative effects (San Felipe et al., 2024).

In this context, the objective of this study was to evaluate the impact of ELS induced by MSEW on the emotional behaviour of male and female mice, as well as its relationship with intestinal permeability and neuroinflammatory markers in the hippocampus. Through a series of behavioural tests, the effects of MSEW on anxiety, depression, and anhedonia were investigated, along with the analysis of intestinal barrier integrity. Additionally, we assessed the expression of inflammation-related proteins in the hippocampus and examined neurogenesis in the hippocampus using immunohistochemistry. These results provide valuable insights into the influence of ELS on mental health and the importance of considering sex as a relevant variable in biomedical research.

## 2. MATERIALS AND METHODS

### Animals

Three male and three female CD1 adult mice aged 10 weeks, used as breeders, were obtained from Charles River (Barcelona, Spain) and housed in our animal facility. The animals were placed in pairs in standard cages and maintained on a 12-hour light/dark cycle (lights on at 8:00 am) with water and food provided ad libitum. Ten days later, the males were removed from the cages. The sex distribution of the offspring was estimated to be 43% males and 57% females. The total number of animals used in this study was 32 mice.

All procedures were performed in accordance with European and Spanish regulations for animal research (Directive 2010/63/EU, Real Decreto 53/2013, and Ley 32/2007) and were approved by the research ethics committees of the University of Málaga (code: 49-2023-A) and the Junta de Andalucía (code: 06/09/2024/127).

### Rearing conditions

The rearing conditions were as previously described (Castro-Zavala et al., 2020, 2021a, 2021b). Newborn mice were randomly assigned to two experimental groups: Standard Nest (SN) and Maternal separation with Early Weaning (MSEW). The day of birth was designated as postnatal day (PD) 0. In the MSEW groups, pups were separated from their mothers for 4 hours per day (9:00 to 13:00) from PD2 to PD5, and for 8 hours per day (9:00 to 17:00) from PD6 to PD16 (**Figure 1A**). During separation, the pups were placed in another cage and room, with their home boxes placed on electric blankets to maintain body temperature until the mothers were returned. Animals in the SN group remained with their mothers until weaning (PD21), while animals in the MSEW group were weaned at PD17. In both groups (SN and MSEW), the cages were cleaned on PD10. Each litter was distributed randomly across the experimental groups to avoid litter effects. For the behavioural tests, the groups were: SN males n=8, SN females n=8, MSEW males n=8 and MSEW females n=8. MSEW protocol does not affect body weight (Gracia-Rubio et al., 2016a; Portero-Tresserra et al., 2018), mortality (George et al., 2010), morbidity (George et al., 2010) or the male/female ratio (Koob and Zorrilla, 2010).

**Figure 1.**
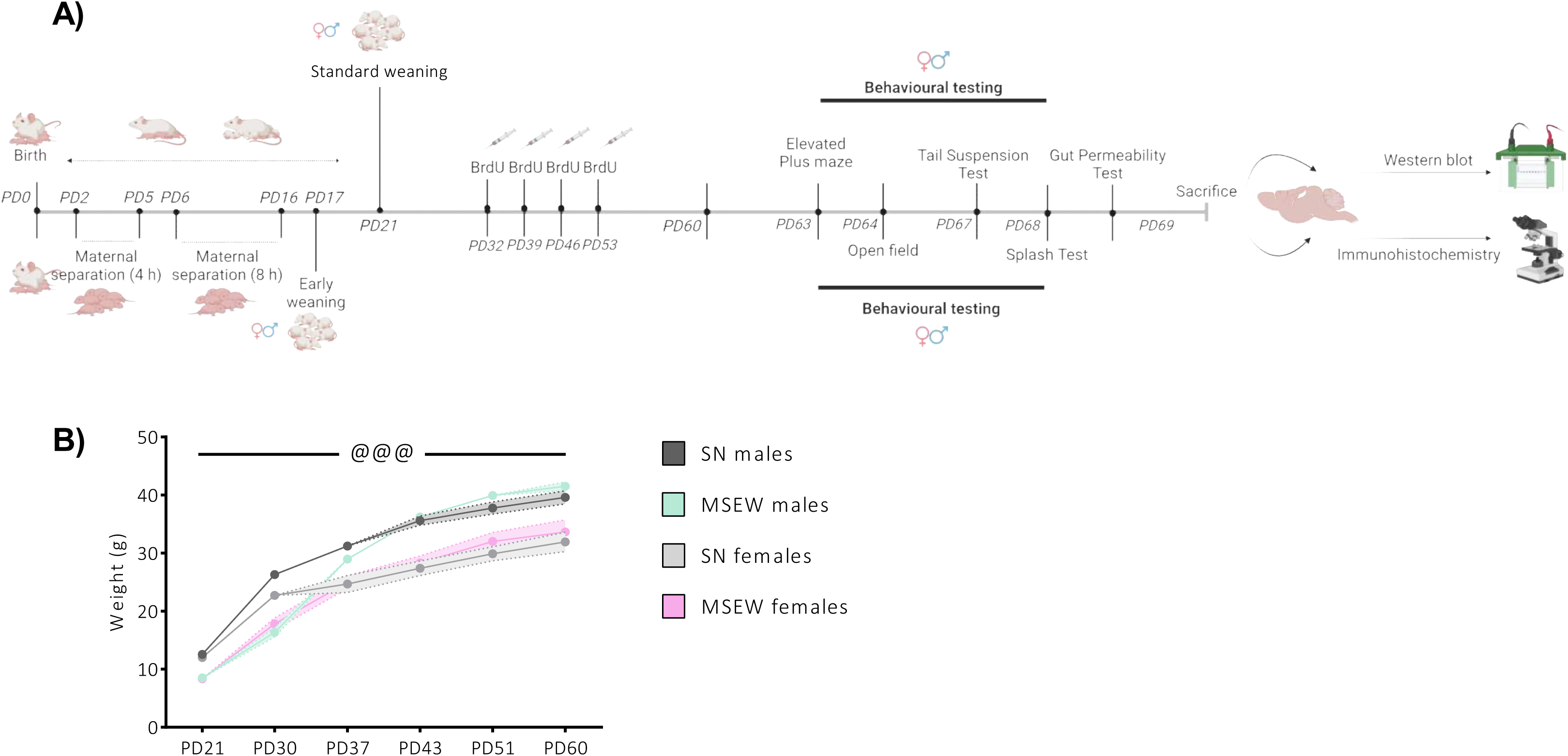
(A) Schematic representation of the MSEW model and the timeline for the experiments (Created in BioRender.com). (B) Mean body weight of male and female mice across different postnatal days. *Sex* main effect of the ANOVA (@@@p<0.001). Data are expressed as mean ± SEM (n=8 per group).

### Bromodeoxyuridine administration

To study adult hippocampal neurogenesis (AHN), bromodeoxyuridine (BrdU, Sigma-Aldrich, Madrid, Spain) was administered weekly on PD 32, 39, 46, and 53 to tag newly generated cells. Mice received two daily intraperitoneal injections of BrdU at a dosage of 75 mg/kg, diluted in physiological saline, with a 4-hour interval between each injection (Mañas-Padilla et al., 2023) (**Figure 1A**).

### Behavioural testing

Behavioural testing began on postnatal day (PD) 63 (**Figure 1A**) . Mice were transported to a noise-isolated room (illuminated at 20 lux) at 8:30 a.m., where they were habituated for at least 30 minutes before starting the behavioural assessment. Sessions were recorded, and spatio-temporal parameters were analysed using Ethovision XT.17 software (Noldus, Wageningen, The Netherlands). Observational scoring was performed by a trained observer who was blind to the mice’s sex and experimental condition, with no prior assumptions about the study’s outcome. The behavioural evaluation was conducted following the same methodology as in previous studies (García-Baos et al., 2022; Melgar-Locatelli et al., 2024a, 2024b). The behavioural schedule was structured as shown in **Figure 1A**.

#### **a)** Elevated Plus Maze (EPM)

The plus-shaped apparatus was positioned at a height of 47 cm from the floor and comprised two unprotected open arms and two enclosed arms (each measuring 29.5 x 5 cm) connected by a central platform (5 x 5 cm). The mouse was introduced (PD 63) onto the central platform and allowed to freely explore the apparatus for a duration of 6 min. Total distance travelled (cm), time spent in the open arms (s) and percentage of distance travelled in the open arms (%) *(% distance in open arms= (distance in open arms*100)/total* distance travelled) were analysed.

#### **b)** Open Field Test (OFT)

On PD 64, mice were placed in the centre of an empty open field (40 x 40, 40 cm high) and allowed to freely explore for 5 min (habituation session). Total distance travelled (cm), time spent (s) in the centre zone (comprising an imaginary central square of 20 x 20 cm) and centre entry frequency were evaluated.

#### **c)** Tail Suspension Test (TST)

A computerised instrument (Bioseb, Bordeaux, France) was used to perform the TST. On PD 67, mice were attached to a hook that was attached to a strain gauge by adhesive tape. Over the course of a 6-minute test, the gauge transmitted motions to a computer, which computed the total immobility time (s) and the percentage of immobility (%).

#### **d)** Splash Test

On PD 68, a splash test was conducted in a standard cage. After five minutes of habituation, mice were placed in a corner of the cage and splashed twice (∼2 × 0.6 ml) with 10% sucrose solution diluted in tap water (Garcia-Mompo et al., 2020; García-Baos et al., 2022). The total time of self-grooming behaviour (licking, stroking and scratching) was manually recorded for 5 minutes.

### Gut permeability assay

Gut permeability was assessed on PD 69, following protocol described by Botía-Sánchez et al (Botía-Sánchez et al., 2023). Briefly, mice were fasted (without access to food and water) for 4 hours. They were then administered fluorescein isothiocyanate (FITC)-coupled dextran 4 kDa (Sigma) at 250 mg/kg via gavage dissolved in 1x PBS.

After 3 hours, mice were deeply anesthetized with intraperitoneal sodium pentobarbital (50 mg/kg body weight), and blood was collected intracardially. Fluorescence intensity was measured in serum samples diluted 1:4 in PBS using a plate reader (Infinite 200Pro) with excitation and emission wavelengths set at 485 nm and 528 nm, respectively. A total of 5 animals per group were used (SN males n=5, SN females n=5, MSEW males n=5, MSEW females n=5), chosen randomly. The autofluorescence emission value of plasma from untreated mice was subtracted from the experimental samples.

### Brain samples collection

After blood collection, mice were intracardially perfused with 0.1 M phosphate buffered saline pH 7.4 (PBS) and subsequently sacrificed by decapitation. Brain samples were immediately extracted; the right hemisphere was post-fixed in paraformaldehyde (PFA) during 48 h at 4°C and cut into coronal (45 μm) vibratome sections for immunohistochemistry, while the left hemisphere was directly frozen at −80°C for protein analysis by western blot.

### Western blot

The left hemisphere of the hippocampus was dissected from frozen brain (−80°C), using the Paxinos and Watson mouse brain atlas (Paxinos and Franklin, 2012) as a reference. Brain samples (17-21 mg per sample) were homogenized in 1 ml of cold radioimmunoprecipitation assay buffer lysis buffer (RIPA). The RIPA buffer consisted of 50-mM Tris–HCl pH 7.4, 150-mM NaCl, 0.5% Sodium Deoxycholate, 1-mM Ethylenediaminetetraacetic acid (EDTA), 1% Triton, 0.1% SDS, 1-mM Na3VO4, 1-mM NaF. Additionally, the homogenization buffer was supplemented with a phosphatase (Phosphatase Inhibitor Cocktail Set III, 524527, Millipore, Darmstadt, Germany) and a protease (complete™ Protease Inhibitor Cocktail, 11836145001, Roche, Basel, Switzerland) inhibitor cocktail. Following a 2 h incubation at 4°C, the suspension was centrifugated at 12,000 rpm for 15 min at 4°C. The resulting protein extracts (obtained from the supernatant) were diluted 1:1 in loading buffer (Ditiotreitol [DTT] 2X) and heated for 5 min at 99°C.

We quantified protein expression levels of TNF-α, NF-kB and BDNF in the hippocampus samples. Tissue protein samples (10–15 μg) were subjected to electrophoresis on 4–12% Criterion XT Precast Bis-Tris gels (3450125, Bio-Rad, California, USA) for 30 min at 80 V, followed by 2 h at 150 V. The separated proteins were then transferred onto a 0.2-μm nitrocellulose membrane (Bio-Rad, USA) using wet transfer equipment (Bio-Rad, USA) for 1 h at 80 V. Ponceau Red staining (10x diluted to 1x in H_2_O) was utilized for protein visualization. Subsequently, the membrane was washed with TBST 1X Tween 20 (150-mM NaCl, 10-mM Tris-HCl, 0.1% Tween 20, pH 7.6) until it became clean and clear. The membrane was blocked with 2% bovine serum albumin-Tris buffered saline Tween 20 (BSA-TBST1X) on a shaker platform at room temperature for 1 h. Next, the membrane was incubated overnight at 4°C with the respective primary antibody (Table 1), diluted in 2% BSA-TBST1X. The following day, the membrane was washed three times for 10 min with TBST 1X and then incubated with an appropriate Horseradish Peroxidase conjugated secondary antibody (Table 1) were diluted 1:10,000 in 2% BSA-TBST 1X for 1 h at room temperature on a shaker. After washing the membrane, it was exposed to a chemiluminescent reagent (Santa Cruz Biotechnology) for 5 min. If required, stripping/reproving steps were performed. The protein bands on the membrane were visualized using chemiluminescence (ChemiDoc Imaging System, Bio-Rad, California, USA) and quantified using ImageJ software (densitometric analysis http://imagej.nih.gov/ij). Normalization was accomplished by using a reference protein, γ-adaptin (Table 1), which was present on the same membrane. The results were expressed as the ratio between the total protein expression and γ-adaptin as described (Bass et al., 2017). A total of 4-6 animals per group were used, chosen randomly. Data was normalized to the SN males’ group.

**Table 1.** Antibodies

| **Antibody** | **# Catalogue** | **RRIDs** | **Dilution** | **Company** |
| --- | --- | --- | --- | --- |
| γ-Adaptin | 610385 | AB_397768 | 1:2000 | BD Biosciences |
| TNF-α | 3707 | AB_2240625 | 1:1000 | Cell signaling |
| NF-kB | 8242 | AB_10859369 | 1:1000 | Cell signaling |
| BDNF | AB1534 | AB_90746 | 1:1000 | Millipore |
| Goat anti-rabbit IgG | W4011 | AB_430833 | 1:10,000 | Promega |
| Goat anti-mouse IgG | W4021 | AB_430834 | 1:10,000 | Promega |

### Immunohistochemistry and cell quantification

Following a 48-hour post-fixation period, the hippocampus of the right hemisphere was sectioned into 45 μm coronal sections, resulting in six equivalent tissue series using a Leica VT1000S vibratome. For free-floating immunohistochemistry, the following steps were undertaken: first, sections were subjected to an antigen retrieval method using EnVision Flex high pH solution (Dako, Glostrup, Denmark) for 1 min in a microwave. Subsequently, an endogenous peroxidase blocking solution consisting of 80% PBS, 10% methanol, and 10% hydrogen peroxide was applied in darkness for 30 min. After PBS rinses, the sections were incubated overnight in the primary antibody, which was diluted in a solution of PBS, 0.5% Triton X-100, and donkey serum. The primary antibody used was rat anti-BrdU (1:500, ab6326, Abcam). On the following day, appropriate biotin-conjugated secondary antibody was incubated for 90 min (rabbit anti-rat, 1:200, 31834, Invitrogen, Carlsbad, USA). The staining process was carried out using the biotin and peroxidase-conjugated extravidin method, employing diaminobenzidine (DAB) and hydrogen peroxide as the chromogen/substrate. PBS rinses followed each step of the protocol.

The dentate gyrus within the dorsal hippocampus (bregma −1.06 mm to −3.08 mm) (Paxinos and Franklin, 2012) as examined for immunohistochemical expression to determine the presence of mentioned specific markers related to AHN. To quantify the cells stained with DAB, detailed photographs of every sixth hippocampal section were captured using an Olympus BX41TF-5 microscope equipped with an Olympus DP70 digital camera (Olympus, Glostrup, Denmark). The software ImageJ (National Institutes of Health, Maryland, USA) was used to measure and analyse the drawn regions of interest. The number of positive cells within each region was counted and expressed as the number of cells per mm2. All immunohistochemical procedures and the cell counting were performed following protocols previously used by our research group (Mañas-Padilla et al., 2023; Melgar-Locatelli et al., 2024b, 2024a).

### Statistical Analysis

The data were assessed for normality (Kolmogorov-Smirnov’s test), sphericity (Mauchly’s test) and homoscedasticity (Levene’s test). A two-way ANOVA was employed with *rearing* and *sex* as independent factors. Post-hoc analysis using the Bonferroni test was conducted when the F statistic achieved p<0.05, indicating a significant main effect and/or interaction. All possible pairwise comparisons were evaluated. For correlation analysis, Pearson correlation coefficients were calculated. Statistical analyses were performed using SPSS Statistics v25. Data are presented as mean ± SEM, and significance was set at p<0.05.

## 3. RESULTS

### MSEW Did Not Affect Animal Weight Over Time, but There Was a Difference Between Males and Females

To confirm whether MSEW affected the weight and growth of the animals, we weighed the mice over time (PD21, PD37, PD43, PD51 and PD60) (Figure 1B). A two-way repeated measures ANOVA revealed a main effect of *sex* (F_1,28_=33.93, p<0.001), with males weighing more than females.

### MSEW Induces Anxiety-like behaviour in Both Sexes, with a Greater Effect on Females in the EPM

We evaluated the effect of MSEW in the EPM (Figure 2). First, we measured the total distance travelled, which showed no significant differences between the groups. For the time spent in open arms and the percentage of distance travelled in open arms, the two-way ANOVA showed a main effect of *rearing* for total time in open arms (F_1,28_=19.21, p<0.001) and percentage of distance travelled in open arms (F_1,28_=10.83, p<0.01). For both total time in open arms and the percentage of distance travelled in open arms, there was an interaction between the factors (F_1,28_=5.65, p<0.05; F_1,28_=7.49, p<0.05, respectively). Bonferroni post-hoc test for total time in open arms revealed that SN males spent more time in the open arms than MSEW females (p<0.05), and SN females spent more time than both MSEW males (p<0.05) and MSEW females (p<0.05). Bonferroni post-hoc tests for the percentage of total distance in open arms showed that SN males spent a higher percentage of time in open arms than MSEW females (p<0.05), and SN females spent a higher percentage of time in open arms than MSEW females (p<0.01).

**Figure 2.**
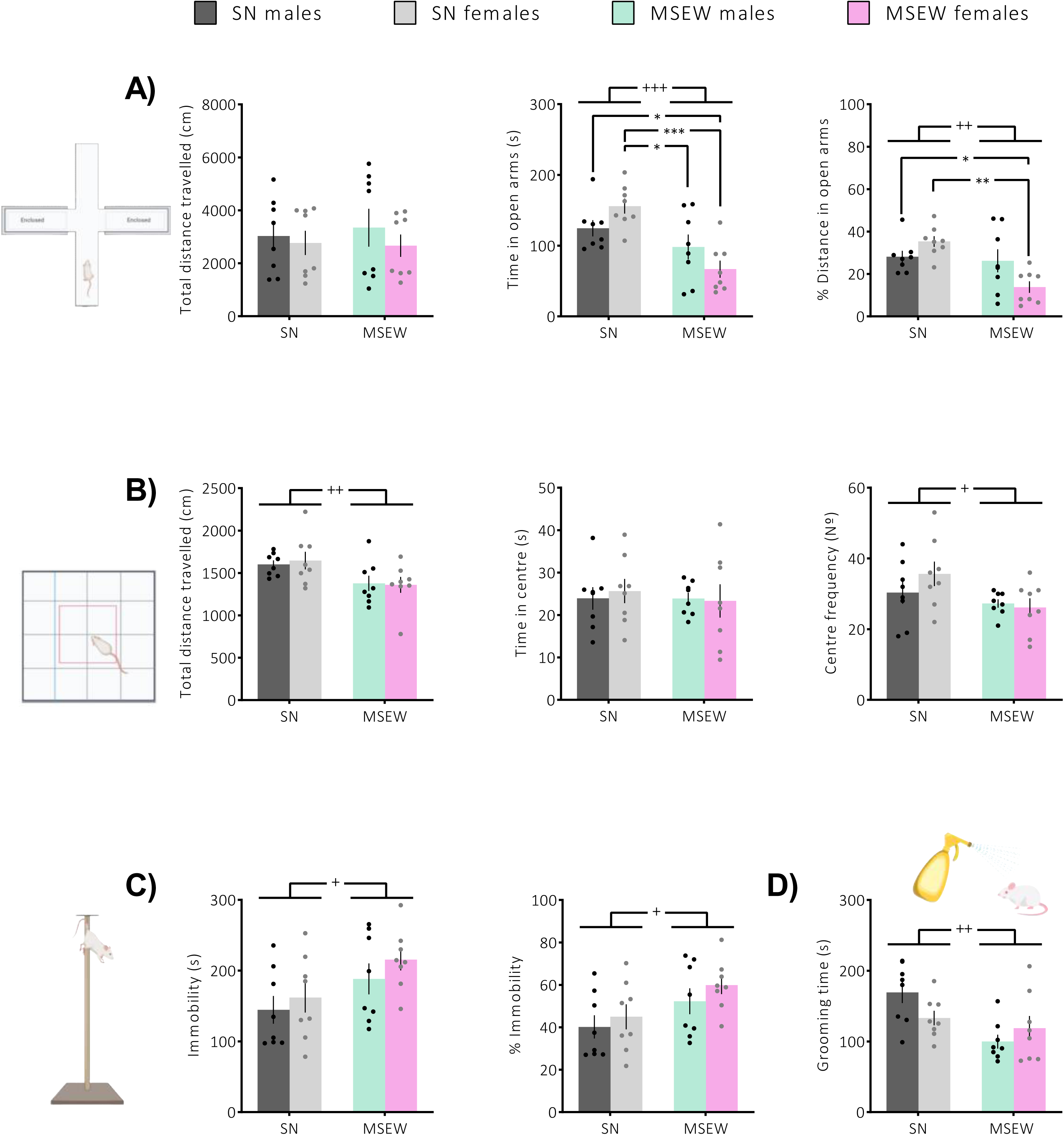
Effects of MSEW on anxiety-like and despair-like behaviours. (A) Total distance travelled, time spent in open arms and percentage of distance in open arms measured in the EPM test. (B) Total distance travelled, time spent in the centre and frequency of centre entries obtained from the open field test. (C) Total immobility time, percentage of immobility in the tail suspension test, and (D) total grooming time in the splash test. *Rearing* main effect of the ANOVA (+p<0.05, ++p<0.01, +++p<0.001). Bonferroni post-hoc comparisons for the interaction *sex × rearing* are indicated with lines (*p<0.05, **p<0.01, ***p<0.001). Data are expressed as mean ± SEM (n=8 per group).

### MSEW Mice Show Reduced Centre Entries Compared to SN Mice in the OFT

We also evaluated the anxiogenic effect induced by the MSEW in the OFT (Figure 2B). The two-way ANOVA for total distance travelled in the OFT showed a main effect of *rearing* (F_1,28_=8.75, p<0.01), with MSEW animals travelling less distance than SN mice. For the time spent in the centre, no significant differences were observed. Regarding the frequency of centre entries, a two-way ANOVA revealed a main effect of *rearing* (F_1,28_=5.29, p<0.05), with MSEW entering the centre less frequently than SN mice.

### Increased Despair-Like Behaviour and Anhedonia in MSEW Mice

The despair-like behaviour was evaluated using the TST (Figure 2C). The two-way ANOVA showed a main effect of *rearing* indicating that MSEW increased total immobility time (F_1,28_=6.12, p<0.05) and the percentage of immobility (F_1,28_=6.12, p<0.05). Additionally, self-care and hedonic behaviour were assessed using the splash test (Figure 2D). The two-way ANOVA indicated that mice exposed to MSEW exhibited decreased self-grooming behaviour when compared to SN mice (F_1,28_=9.73, p<0.01).

### MSEW Increases Gut Permeability and Correlates with Anxiety-Like Behaviour and Anhedonia

We assessed whether MSEW alters gut barrier integrity using the FITC-dextran method (Figure 3A). A two-way ANOVA revealed a main effect of *rearing* on gut permeability (F_1,16_=36.63, p<0.001) indicating that MSEW mice showed higher permeability.

**Figure 3.**
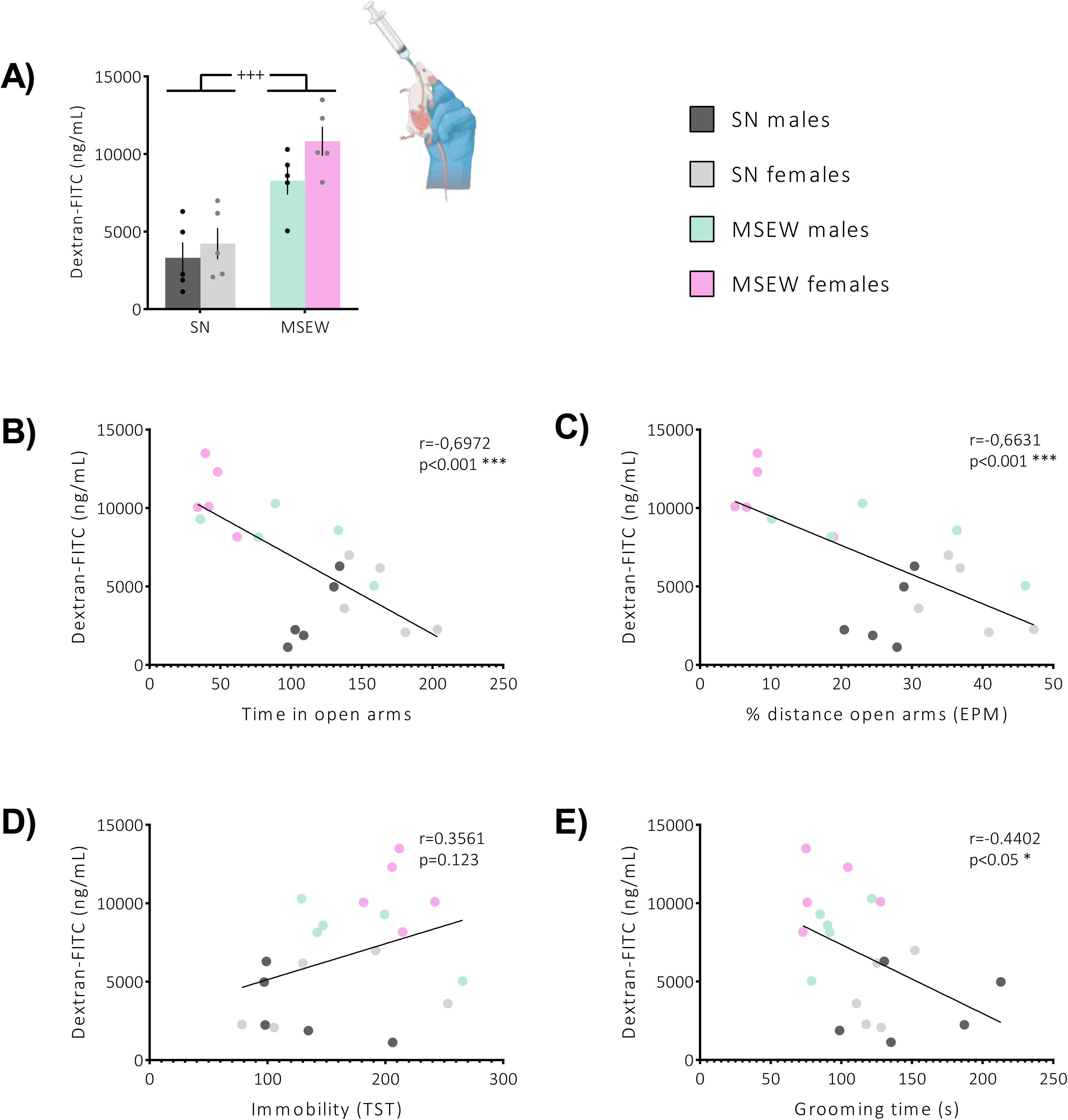
Effects of MSEW on gut permeability and correlation with behavioural results. (A) Gut permeability *in vivo* with FITC-dextran. Data are expressed as mean ± SEM (n=5 per group). A scatter blot illustrating the correlation between gut permeability and (B) time in the open arms of the EPM, (C) percentage of distance in the open arms of the EPM, (D) total immobility time in the TST, and (E) total grooming time in the splash test (n=5 per group, run in duplicate).

Additionally, we calculate the correlation between gut permeability and several behavioural measures: time in open arms (Figure 3B), % distance travelled in open arms (Figure 3C), immobility time (Figure 3D) and grooming time (Figure 3E). There was a significant and negative correlation between gut permeability and time spent in open arms (r=-0.697, p<0.001, n=20), % distance travelled in open arms (r=-0.663, p<0.001, n=20) and grooming time (r=-0.440, p<0.05, n=20).

Linear regression analysis suggests that time spent in open arms (F_1,18_=14.12, p<0.001) (y=-186.1x+11347), % distance travelled in open arms (F_1,18_=17.03, p<0.001) (y=-49.81x+11936) and grooming time (F_1,18_=4.32, p<0.05) (y=-43.97x+11762), could predict gut permeability in mice.

### Reduced TNF-α but Higher NF-kB Protein Levels in The Hippocampus of MSEW Mice

We investigated whether MSEW induces increased protein expression of specific neuroinflammatory markers in the hippocampus. A two-way ANOVA for NF-kB (Figure 4A) protein levels in the hippocampus showed a main effect of *rearing* (F_1,20_=6.52, p<0.05) and a main effect of *sex* (F_1,20_=27.75, p<0.001), indicating higher NF-kB levels in both MSEW mice and in males. In contrast, a two-way ANOVA for TNF-α (Figure 4B) revealed a main effect of *rearing* (F_1,20_ = 6.63, p<0.05) and an interaction between *rearing* and *sex* (F_1,20_ = 5.41, p<0.05). The main effect of *rearing* indicated reduced TNF-α levels in MSEW mice compared to SN mice. Bonferroni post-hoc analysis for the interaction showed this reduction was particularly significant in MSEW male mice (p<0.05).

**Figure 4.**
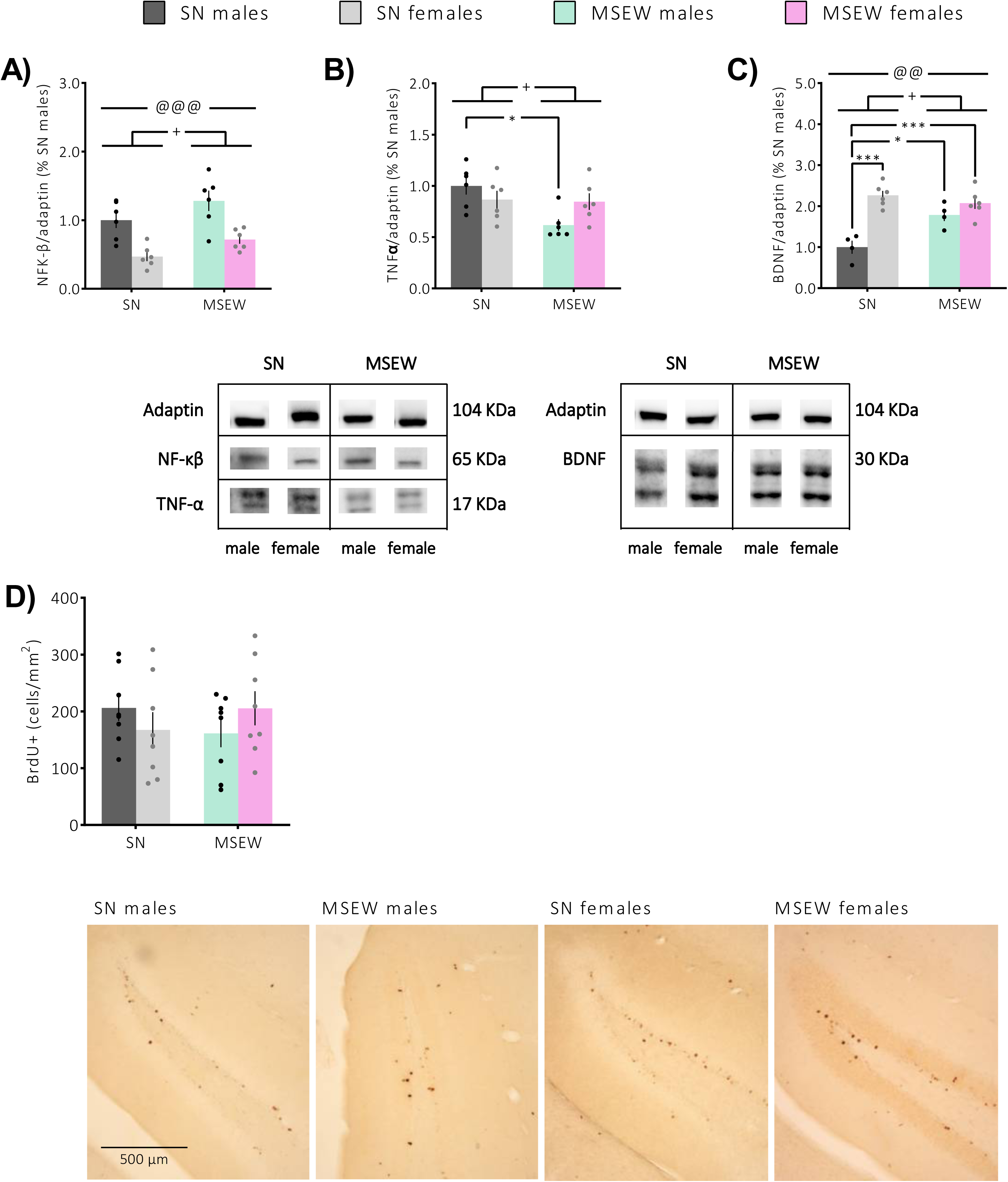
Protein expression of (A) TNFα, (B) NF-kB and (C) BDNF in the hippocampus (n=4-6 per group). (D) Number of BrdU+ cells that survived until the end of the experiment (n=8 per group). *Sex* main effect of the ANOVA (@@p<0.01, @@@p<0.001). *Rearing* main effect of the ANOVA (+p<0.05). Bonferroni post-hoc comparisons for the interaction *sex × rearing* are indicated with lines (*p<0.05, ***p<0.001). Data are expressed as mean ± SEM.

### MSEW Increases BDNF Expression in the Hippocampus, Particularly in Male Mice

We also assessed BDNF protein expression in the Hippocampus (Figure 4C). A two-way ANOVA revealed main effects of *rearing* (F_1,16_=4.50, p<0.05) and *sex* (F_1,16_=30.59, p<0.001), as well as a significant interaction between these factors (F_1,16_=12.08, p<0.01). These results indicated that females have higher BDNF expression in the hippocampus and that MSEW mice express higher BDNF than SN mice. Bonferroni post-hoc analysis for the interaction showed that SN females exhibited higher BDNF levels than SN males (p<0.001). Additionally, MSEW increased BDNF expression particularly in males (p<0.05) and MSEW females showed higher BDNF levels than SN males (p<0.001).

### MSEW Does Not Alter the Survival of Newly-generated Hippocampal Cells

We investigated whether MSEW affects adult hippocampal neurogenesis (AHN) in the hippocampus (Figure 4D). A two-way ANOVA revealed no significant effects. This indicates that there was no alteration in the number of BrdU+ cells that survived until the end of the experiment.

## DISCUSSION

This study demonstrates that MSEW induces a lasting increase in anxiety-like behaviour, particularly in females, as well as an increase in depression-like behaviours and anhedonia in mice of both sexes. Regarding neuroinflammation markers, our analysis revealed that MSEW increased NF-kB expression in the hippocampus and reduced TNF-α expression in the same region, particularly in males. Additionally, when measuring BDNF levels in the hippocampus, we found that MSEW increased BDNF expression in males, although females showed a higher baseline expression in this region compared to males. Our results also showed no significative changes in adult hippocampal neurogenesis (cell survival). Furthermore, MSEW led to an increase in intestinal permeability. We observed a negative correlation between intestinal permeability and both the time spent in the open arms and the distance travelled in these arms in the EPM. Conversely, we found a positive correlation between intestinal permeability and the immobility time in the TST and grooming time in the splash test. These results suggest that increased intestinal permeability is associated with heightened anxiety- and depression-like behaviours. Importantly, these changes in intestinal permeability were not transient, but rather long-lasting, reinforcing the idea that the impact of MSEW on gut health has enduring consequences for emotional and behavioural regulation.

Previous studies using maternal separation, have demonstrated similar increases in anxiety- (Millstein and Holmes, 2007; Kestering-Ferreira et al., 2021; Veenit et al., 2021; De Santa et al., 2024) and depression-like behaviours (Millstein and Holmes, 2007; Récamier-Carballo et al., 2017; Castro-Zavala et al., 2021b; Veenit et al., 2021; De Santa et al., 2024), supporting our findings. However, it is important to note that most of these studies have focused solely on male subjects, leaving the effects observed in females largely unexplored, including potential sex-related variations. In our study, by employing both sexes, we have been able to evaluate the differential effects between males and females. A previous study reported that female mice exposed to a maternal separation protocol spent less time in the open arms compared to the control group, whereas males did not show this effect (Kong et al., 2023). Our results align with the hypothesis that maternal separation leads to increased anxiety-like behaviour, particularly in females, as this sex displayed a decrease in time spent and percentage of distance travelled in the open arms during the EPM. Like the previous study (Kong et al., 2023), we found more pronounced anxiety-like behaviours in females exposed to ELS in the EPM. This difference may be attributed to hormonal regulation of stress responses, as females typically exhibit greater activation of the hypothalamic-pituitary-adrenal (HPA) axis following stress (Goel et al., 2014; Leistner and Menke, 2020; Veenit et al., 2021; Kong et al., 2023).

Although we expected similar results in the OFT, we observed a decrease in entries into the centre in animals exposed to MSEW, but no sex differences. This discrepancy between the two tests, can be explained by the fact that they are different tests and contexts for measuring anxiety. First, both tests assess different components of anxiety; the EPM focuses on behaviour related to exposure to an elevated and unprotected environment, while the OFT assesses exploration in a larger, less structured space, which may influence how animals express their anxiety (Ramos et al., 2008; Mañas-Padilla et al., 2021; Ronquillo et al., 2023). Additionally, the EPM was conducted on the first day, when the animals were less habituated to the experimental environment, which could have increased their anxiety due to the novelty of the space and the presence of both open and closed, unprotected areas. In contrast, the OFT, performed the following day, took place in a more familiar environment for the animals, which may have reduced their anxiety levels, as the lack of clear threat cues in the open field could have encouraged more exploratory behaviour without eliciting significant anxiety responses. However, the fact that the effect of MSEW persisted indicates that this factor is more potent, continuing to show a greater tendency toward anxiety compared to the control group.

In addition, we evaluated depression-like responses. In a previous study that employed the same MSEW model to evaluate its effect on depression-like behaviour using the TST in CD1 mice, both females and males were examined (Castro-Zavala et al., 2021b). It was observed that MSEW generally increased depression-related behaviours, although males were particularly affected, showing a greater duration of immobility (Castro-Zavala et al., 2021b). In the present study, we found that MSEW leads to a general increase in immobility in the TST, which is consistent with previous findings (Castro-Zavala et al., 2021b) . However, we did not observe sex differences in our results. This discrepancy may be explained by differences in methodology: while our evaluation of the TST was conducted automatically, the other study used manual assessment. Additionally, in our protocol, animals had undergone several behavioural tests prior to the TST, unlike in the previous study, which could influence previous experiences and habituation. Moreover, the automatic assessment method alters the criteria for mobility and immobility detection, as it relies on predefined movement thresholds rather than manual scoring. Another depression-related behaviour is anhedonia, which was measured using the splash test. We observed a decrease in grooming time in MSEW mice, regardless of sex. These results are consistent with previous findings (Amiri et al., 2016; Al-Shudifat et al., 2024; Rostami-Faradonbeh et al., 2024), which also reported reduced grooming activity in maternally separated mice. However, all these studies evaluated only male mice, whereas in our study, we included both sexes, demonstrating a similar effect in female mice as well. This additional consideration of both male and female mice provides further insight into potential sex differences in response to MSEW, offering a more comprehensive understanding of the behavioural changes associated with this model.

Despite previous studies evaluating the effects of ELS on mood disorders, our research provides new insights by measuring intestinal permeability induced by MSEW and exploring its potential correlation with the emergence of these mood disorders. This adds support to the gut-brain axis hypothesis (Cryan et al., 2019; Margolis et al., 2021; Socała et al., 2021; Mitrea et al., 2022), which is less frequently explored in the context of ELS. Our findings indicate that MSEW significantly increases gut permeability in the adulthood in both sexes, as demonstrated by the results of the FITC-dextran method.

Consistently, Tao et al. (2023) observed that maternal separation increased intestinal permeability, as evidenced by both the FITC-dextran method and microscopic examination of intestinal tissue, which revealed weakened epithelial tight junctions. They also found that these changes persisted from adolescence into adulthood, with greater intestinal permeability observed in adulthood. These findings align with the gut-brain axis hypothesis, which posits that gut health can significantly influence brain function and emotional states (Cryan et al., 2019; Margolis et al., 2021; Socała et al., 2021; Mitrea et al., 2022). Therefore, this increase in gut permeability may play a crucial role in the behavioural changes observed in our study. We observed that altered intestinal permeability correlated with depression-like behaviours, as well as with reduced time spent in the open arms and lower percentage of distance travelled in these arms, suggesting also a link between increased gut permeability and anxiety-like behaviours.

The correlation we found between gut permeability and grooming behaviour suggests that alterations in gut integrity may contribute to anhedonia, reflecting a broader impact of gut health on emotional well-being. A study on chronic restraint stress in mice reported increased anxiety- and depression-like behaviours alongside intestinal alterations, including changes in neurotransmitter levels, enzyme activity, and barrier integrity (Deng et al., 2021). These changes may underlie the heightened vulnerability to stress-induced mood disorders. Moreover, another study also reported a negative correlation between biomarkers of gut barrier integrity and anxiety-like behaviour, suggesting that greater intestinal damage is associated with more pronounced anxiety-like behaviour (Cai et al., 2024). Furthermore, previous studies have demonstrated that interventions targeting gut permeability can mitigate anxiety-like behaviors. For instance, the administration of resveratrol was found to reduce maternal separation-induced anxiety-like behavior in male mice; however, this study was limited to male subjects and assessed animals in late adulthood (postnatal day 90) (Wei et al., 2023). In contrast, a study that included both sexes found that oleanolic acid and ursolic acid reversed the anxiety-related effects of maternal separation specifically in female mice, as maternal separation did not induce anxiety-like behavior in males (Kong et al., 2023). Additionally, these compounds also reversed anhedonic-like behavior, as measured by the splash test, highlighting their potential role in modulating emotional responses through gut health restoration (Kong et al., 2023). Our study provides a novel contribution by showing that MSEW not only induces both anxiety and depression-like behaviors but also correlates with increased intestinal permeability. Importantly, we assessed both male and female mice, which allowed us to demonstrate that the MSEW-induced effects on anxiety and depression were present in both sexes, providing a more comprehensive understanding of how gut health alterations may impact emotional regulation across genders. This highlights the potential relevance of targeting gut permeability as a therapeutic approach for both male and female subjects, addressing the emotional and behavioral dysregulation associated with MSEW.

It has been proposed that alterations in the intestinal barrier may contribute to anxiety and depression by triggering neuroinflammation through the passage of endotoxins into the bloodstream(De Punder and Pruimboom, 2015). Over time, intestinal changes can lead to systemic inflammation, manifesting as neuroinflammation (Thevaranjan et al., 2017; Haruwaka et al., 2019), which can affect synaptic plasticity (Doan et al., 2015; Lima Giacobbo et al., 2019). Accordingly, we evaluated neuroinflammatory markers using western blot analysis to assess TNF-α and NF-κB levels in the hippocampus, along with the expression of the neurotrophin BDNF.

Our results revealed baseline sex differences in BDNF expression, with females exhibiting higher levels in the hippocampus compared to males. To our knowledge, there are not many studies evaluating BDNF expression in mice while considering sex differences. However, our results are consistent with recent findings showing that BDNF mRNA levels in female mice were significantly higher than those in male mice at 3 months of age, which is similar to the age of the mice in the present study (Matsuoka et al., 2024). This elevation could be due to the effects of estrogen on BDNF expression (Franklin and Perrot-Sinal, 2006; Sohrabji and Lewis, 2006), as estrogen treatment has been observed to increase BDNF levels (Bimonte-Nelson et al., 2004).

Upon exposure to MSEW, both sexes showed increased NF-κB protein levels, which aligns with previous findings linking ELS to neuroinflammatory responses (Gracia-Rubio et al., 2016b; San Felipe et al., 2024). This increase is expected, as NF-κB plays a crucial role in microglial activation and inflammatory signalling suggesting a shared inflammatory response to MSEW (Dresselhaus and Meffert, 2019). However, only MSEW male mice exhibited changes in TNF-α and BDNF levels suggesting a sex-specific regulatory mechanism in response to chronic stress. In males, the reduction in TNF-α alongside increased NF-κB may indicate a shift toward non-canonical NF-κB activation. While TNF-α typically stimulates NF-κB via the canonical pathway (Lawrence, 2009), its reduction suggests an alternative activation mechanism through cytokines such as LTβ, CD40L, BAFF, and RANKL (Yu et al., 2020). This pathway may contribute to a more sustained but potentially regulated inflammatory response

In addition, the male-specific increase in BDNF in MSEW mice suggests a potential compensatory mechanism aimed at mitigating stress-induced neural damage. NF-κB has been implicated in synaptic plasticity and neuronal repair by inducing BDNF expression (Sochocka et al., 2016; Dresselhaus and Meffert, 2019; Zaghloul et al., 2020).

Based on previous literature, we expected that MSEW would reduce adult hippocampal neurogenesis (AHN), given its known vulnerability to early-life stress (ELS) and its role in modulating emotional behaviors (Hulshof et al., 2011; Lajud et al., 2012). AHN is closely regulated by BDNF and is often compromised under neuroinflammatory conditions (Kuipers et al., 2016; Sung et al., 2020; Salta et al., 2023; Khalil, 2024; Khoury et al., 2024). The observed increase in BDNF, despite unchanged AHN levels, suggests that this neurotrophic factor may help preserve neurogenesis, counteracting the potential negative impact of neuroinflammation.

In contrast, females, with already higher baseline BDNF levels, may rely on distinct regulatory mechanisms to buffer the effects of ELS without requiring further upregulation of this neurotrophic factor. Their response to MSEW suggests that while NF-κB activation occurs similarly to males, its downstream effects on TNF-α and BDNF are different, potentially due to sex-specific hormonal or epigenetic influences. These findings underscore the importance of sex-specific pathways in stress-induced neuroinflammation and neuronal adaptation. While both sexes exhibit NF-κB activation in response to MSEW, males appear to engage additional compensatory mechanisms involving TNF-α downregulation and BDNF upregulation, which may represent an adaptive strategy to counteract prolonged stress effects on brain function and behavior.

Taken together, these findings contribute to the growing body of literature suggesting that early life stressors, such as MSEW, induce long-term alterations in gut barrier function, which, in turn, correlate with emotional and behavioral dysregulation. Most importantly, our study highlights intestinal permeability as a biomarker for future research into the mechanisms underlying mood disorders, particularly in contexts of chronic and prolonged stress. Given the observed correlation between increased gut permeability and anxiety- and depression-like behaviors, targeting gut barrier integrity may represent a novel therapeutic avenue for mitigating stress-induced emotional dysfunction. Future studies should explore whether interventions aimed at restoring gut integrity could counteract the neuroinflammatory and behavioural effects of ELS in both sexes.

## AUTHOR CONTRIBUTIONS

**Adriana Castro-Zavala:** conceptualization, methodology, formal analysis, investigation, writing-original draft preparation, visualization, funding acquisition, project administration, supervision. **Ana E. Nieto-Nieves**: methodology, investigation, visualization. **Sonia Melgar-Locatelli**: methodology, investigation, visualization. **M. Carmen Mañas-Padilla:** methodology, investigation, visualization. **Lidia Medina-Rodríguez**: methodology, investigation. **Estela Castilla-Ortega**: conceptualization, methodology, formal analysis, investigation, writing-original draft preparation, visualization, funding acquisition, project administration, supervision.

## AKNOWLEDGEMENTS

## Acknowledgements

This study was funded by Grant PID2020-114374RB-I00 from MCIN/AEI/10.13039/501100011033 (to E.C-O.), Universidad de Málaga (B.1. Ayudas para proyectos dirigidos por jóvenes investigadores B1-2022_05 to A.C.-Z,), and Universidad de Málaga (C.2. II Plan Propio de Investigación, Transferencia y Divulgación Científica). Funding for open access charge: Universidad de Málaga / CBUA.

A.C.Z. holds a postdoctoral research contract from the Secretaría General de Universidades, Investigación y Tecnología–Junta de Andalucía (POSTDOC21_00365) and a Sara Borrell contract from the Instituto de Salud Carlos III (CD24/00041). A.N.-N. holds a predoctoral grant from the Spanish Ministry of Science, Innovation and Universities (FPU22/02044). The authors acknowledge the IBIMA’s common research support structure—ECAI (Centro de Experimentación y Conducta Animal; University of Malaga)—for the maintenance of the mice. We also thank Ana Gavito Collado for her assistance, collaboration, and guidance in the laboratory.

## REFERENCES

1. Al-Shudifat, A. E., Qnais, E., Bseiso, Y., Wedyan, M., Gammoh, O., Alqudah, M., et al. (2024). Antidepressant potential of β-caryophyllene in maternal separation-induced depression-like in mice: A focus on oxidative stress and nitrite levels. Phytomedicine Plus 4, 100624. doi: 10.1016/J.PHYPLU.2024.100624

2. Amiri, S., Amini-Khoei, H., Mohammadi-Asl, A., Alijanpour, S., Haj-Mirzaian, A., Rahimi-Balaei, M., et al. (2016). Involvement of D1 and D2 dopamine receptors in the antidepressant-like effects of selegiline in maternal separation model of mouse. Physiol Behav 163, 107–114. doi: 10.1016/J.PHYSBEH.2016.04.052

3. Baracz, S. J., Everett, N. A., and Cornish, J. L. (2020). The impact of early life stress on the central oxytocin system and susceptibility for drug addiction: Applicability of oxytocin as a pharmacotherapy. Neurosci Biobehav Rev 110, 114–132. doi: 10.1016/j.neubiorev.2018.08.014

4. Bimonte-Nelson, H. A., Nelson, M. E., and Granholm, A. C. E. (2004). Progesterone counteracts estrogen-induced increases in neurotrophins in the aged female rat brain. Neuroreport 15, 2659–2663. doi: 10.1097/00001756-200412030-00021

5. Björkholm, C., and Monteggia, L. M. (2016). BDNF -A key transducer of antidepressant effects. Neuropharmacology 102, 72–79. doi: 10.1016/j.neuropharm.2015.10.034

6. Boersma, M. C. H., Dresselhaus, E. C., de Biase, L. M., Mihalas, A. B., Bergles, D. E., and Meffert, M. K. (2011). A requirement for nuclear factor-kappaB in developmental and plasticity-associated synaptogenesis. J Neurosci 31, 5414–5425. doi: 10.1523/JNEUROSCI.2456-10.2011

7. Botía-Sánchez, M., Galicia, G., Albaladejo-Marico, L., Toro-Domínguez, D., Morell, M., Marcos-Fernández, R., et al. (2023). Gut epithelial barrier dysfunction in lupus triggers a differential humoral response against gut commensals. Front Immunol 14. doi: 10.3389/FIMMU.2023.1200769/FULL

8. Cai, Y., Deng, W., Yang, Q., Pan, G., Liang, Z., Yang, X., et al. (2024). High-fat diet-induced obesity causes intestinal Th17/Treg imbalance that impairs the intestinal barrier and aggravates anxiety-like behavior in mice. Int Immunopharmacol 130. doi: 10.1016/J.INTIMP.2024.111783

9. Castro-Zavala, A., Martín-Sánchez, A., Luján, M. Á., and Valverde, O. (2021a). Maternal separation increases cocaine intake through a mechanism involving plasticity in glutamate signalling. Addiction biology 26, e12911. doi: 10.1111/adb.12911

10. Castro-Zavala, A., Martín-Sánchez, A., Montalvo-Martínez, L., Camacho-Morales, A., and Valverde, O. (2021b). Cocaine-seeking behaviour is differentially expressed in male and female mice exposed to maternal separation and is associated with alterations in AMPA receptors subunits in the medial prefrontal cortex. Prog Neuropsychopharmacol Biol Psychiatry 109. doi: 10.1016/j.pnpbp.2021.110262

11. Castro-Zavala, A., Martín-Sánchez, A., and Valverde, O. (2020). Sex differences in the vulnerability to cocaine’s addictive effects after early-life stress in mice. Eur Neuropsychopharmacol 32, 12–24. doi: 10.1016/J.EURONEURO.2019.12.112

12. Cryan, J. F., O’riordan, K. J., Cowan, C. S. M., Sandhu, K. V., Bastiaanssen, T. F. S., Boehme, M., et al. (2019). The microbiota-gut-brain axis. Physiol Rev 99, 1877– 2013. doi: 10.1152/PHYSREV.00018.2018

13. De Punder, K., and Pruimboom, L. (2015). Stress Induces Endotoxemia and Low-Grade Inflammation by Increasing Barrier Permeability. Front Immunol 6, 15. doi: 10.3389/FIMMU.2015.00223

14. De Santa, F., Strimpakos, G., Marchetti, N., Gargari, G., Torcinaro, A., Arioli, S., et al. (2024). Effect of a multi-strain probiotic mixture consumption on anxiety and depression symptoms induced in adult mice by postnatal maternal separation. Microbiome 12. doi: 10.1186/S40168-024-01752-W

15. Deng, Y., Zhou, M., Wang, J., Yao, J., Yu, J., Liu, W., et al. (2021). Involvement of the microbiota-gut-brain axis in chronic restraint stress: disturbances of the kynurenine metabolic pathway in both the gut and brain. Gut Microbes 13, 1–16. doi: 10.1080/19490976.2020.1869501

16. DiSabato, D. J., Quan, N., and Godbout, J. P. (2016). Neuroinflammation: The Devil is in the Details. J Neurochem 139, 136. doi: 10.1111/JNC.13607

17. Doan, L., Manders, T., and Wang, J. (2015). Neuroplasticity underlying the comorbidity of pain and depression. Neural Plast 2015, 504691. doi: 10.1155/2015/504691

18. Dresselhaus, E. C., and Meffert, M. K. (2019). Cellular Specificity of NF-κB Function in the Nervous System. Front Immunol 10. doi: 10.3389/FIMMU.2019.01043

19. Frank, D., Zlotnik, A., Kofman, O., Grinshpun, J., Severynovska, O., Brotfain, E., et al. (2019). Early life stress induces submissive behavior in adult rats. Behavioural Brain Research 372. doi: 10.1016/j.bbr.2019.112025

20. Franklin, T. B., and Perrot-Sinal, T. S. (2006). Sex and ovarian steroids modulate brain-derived neurotrophic factor (BDNF) protein levels in rat hippocampus under stressful and non-stressful conditions. Psychoneuroendocrinology 31, 38–48. doi: 10.1016/J.PSYNEUEN.2005.05.008

21. García-Baos, A., Gallego-Landin, I., Ferreres-Álvarez, I., Puig-Reyne, X., Castro-Zavala, A., Valverde, O., et al. (2022). Effects of fast-acting antidepressant drugs on a postpartum depression mice model. Biomedicine & Pharmacotherapy 154, 113598. doi: 10.1016/J.BIOPHA.2022.113598

22. Garcia-Mompo, C., Curto, Y., Carceller, H., Gilabert-Juan, J., Rodriguez-Flores, E., Guirado, R., et al. (2020). Δ-9-Tetrahydrocannabinol treatment during adolescence and alterations in the inhibitory networks of the adult prefrontal cortex in mice subjected to perinatal NMDA receptor antagonist injection and to postweaning social isolation. Translational Psychiatry 2020 10:1 10, 1–13. doi: 10.1038/s41398-020-0853-3

23. George, E. D., Bordner, K. A., Elwafi, H. M., and Simen, A. A. (2010). Maternal separation with early weaning: a novel mouse model of early life neglect. BMC Neurosci 11, 123. doi: 10.1186/1471-2202-11-123

24. Goel, N., Workman, J. L., Lee, T. T., Innala, L., and Viau, V. (2014). Sex differences in the HPA axis. Compr Physiol 4, 1121–1155. doi: 10.1002/CPHY.C130054

25. Gracia-Rubio, I., Martinez-Laorden, E., Moscoso-Castro, M., Milanés, V., Laorden, L., and Valverde, O. (2016a). Maternal Separation Impairs Cocaine-Induced Behavioural Sensitization in Adolescent Mice. PLoS One 11, e0167483. doi: 10.1371/journal.pone.0167483

26. Gracia-Rubio, I., Moscoso-Castro, M., Pozo, O. J., Marcos, J., Nadal, R., and Valverde, O. (2016b). Maternal separation induces neuroinflammation and long-lasting emotional alterations in mice. Prog Neuropsychopharmacol Biol Psychiatry 65, 104–17. doi: 10.1016/j.pnpbp.2015.09.003

27. Guo, B., Zhang, M., Hao, W., Wang, Y., Zhang, T., and Liu, C. (2023). Neuroinflammation mechanisms of neuromodulation therapies for anxiety and depression. Transl Psychiatry 13. doi: 10.1038/S41398-022-02297-Y

28. Haruwaka, K., Ikegami, A., Tachibana, Y., Ohno, N., Konishi, H., Hashimoto, A., et al. (2019). Dual microglia effects on blood brain barrier permeability induced by systemic inflammation. Nat Commun 10. doi: 10.1038/S41467-019-13812-Z

29. Hulshof, H. J., Novati, A., Sgoifo, A., Luiten, P. G. M., Den Boer, J. A., and Meerlo, P. (2011). Maternal separation decreases adult hippocampal cell proliferation and impairs cognitive performance but has little effect on stress sensitivity and anxiety in adult Wistar rats. Behavioural Brain Research 216, 552–560. doi: 10.1016/J.BBR.2010.08.038

30. Imielski, Y., Schwamborn, J. C., Lüningschrör, P., Heimann, P., Holzberg, M., Werner, H., et al. (2012). Regrowing the adult brain: NF-κB controls functional circuit formation and tissue homeostasis in the dentate gyrus. PLoS One 7. doi: 10.1371/JOURNAL.PONE.0030838

31. Jiang, J., Fu, Y., Tang, A., Gao, X., Zhang, D., Shen, Y., et al. (2023). Sex difference in prebiotics on gut and blood–brain barrier dysfunction underlying stress-induced anxiety and depression. CNS Neurosci Ther 29, 115. doi: 10.1111/CNS.14091

32. Kestering-Ferreira, E., Tractenberg, S. G., Lumertz, F. S., Orso, R., Creutzberg, K. C., Wearick-Silva, L. E., et al. (2021). Long-term Effects of Maternal Separation on Anxiety-Like Behavior and Neuroendocrine Parameters in Adult Balb/c Mice. Chronic Stress 5. doi: 10.1177/24705470211067181

33. Khalil, M. H. (2024). The BDNF-Interactive Model for Sustainable Hippocampal Neurogenesis in Humans: Synergistic Effects of Environmentally-Mediated Physical Activity, Cognitive Stimulation, and Mindfulness. International Journal of Molecular Sciences 2024, Vol. 25, Page 12924 25, 12924. doi: 10.3390/IJMS252312924

34. Khoury, Z. S., Sohail, F., Wang, J., Mendoza, M., Raake, M., Silat, M. T., et al. (2024). Neuroinflammation: A Critical Factor in Neurodegenerative Disorders. Cureus 16, e62310. doi: 10.7759/CUREUS.62310

35. Kong, C. H., Park, K., Kim, D. Y., Kim, J. Y., Kang, W. C., Jeon, M., et al. (2023). Effects of oleanolic acid and ursolic acid on depression-like behaviors induced by maternal separation in mice. Eur J Pharmacol 956, 175954. doi: 10.1016/J.EJPHAR.2023.175954

36. Koo, J. W., Russo, S. J., Ferguson, D., Nestler, E. J., and Duman, R. S. (2010). Nuclear factor-κB is a critical mediator of stress-impaired neurogenesis and depressive behavior. Proc Natl Acad Sci U S A 107, 2669. doi: 10.1073/PNAS.0910658107

37. Koob, G. F., and Zorrilla, E. P. (2010). Neurobiological mechanisms of addiction: focus on corticotropin-releasing factor. Curr Opin Investig Drugs 11, 63–71.

38. Kuipers, S. D., Trentani, A., Tiron, A., Mao, X., Kuhl, D., and Bramham, C. R. (2016). BDNF-induced LTP is associated with rapid Arc/Arg3.1-dependent enhancement in adult hippocampal neurogenesis. Scientific Reports 2016 6:1 6, 1–14. doi: 10.1038/srep21222

39. Lajud, N., Roque, A., Cajero, M., Gutiérrez-Ospina, G., and Torner, L. (2012). Periodic maternal separation decreases hippocampal neurogenesis without affecting basal corticosterone during the stress hyporesponsive period, but alters HPA axis and coping behavior in adulthood. Psychoneuroendocrinology 37, 410–420. doi: 10.1016/J.PSYNEUEN.2011.07.011

40. Lawrence, T. (2009). The Nuclear Factor NF-κB Pathway in Inflammation. Cold Spring Harb Perspect Biol 1. doi: 10.1101/CSHPERSPECT.A001651

41. Leistner, C., and Menke, A. (2020). Hypothalamic–pituitary–adrenal axis and stress. Handb Clin Neurol 175, 55–64. doi: 10.1016/B978-0-444-64123-6.00004-7

42. Lima Giacobbo, B., Doorduin, J., Klein, H. C., Dierckx, R. A. J. O., Bromberg, E., and de Vries, E. F. J. (2019). Brain-Derived Neurotrophic Factor in Brain Disorders: Focus on Neuroinflammation. Mol Neurobiol 56, 3295–3312. doi: 10.1007/S12035-018-1283-6

43. Mañas-Padilla, M. C., Ávila-Gámiz, F., Gil-Rodríguez, S., Ladrón de Guevara-Miranda, D., Rodríguez de Fonseca, F., Santín, L. J., et al. (2021). Persistent changes in exploration and hyperactivity coexist with cognitive impairment in mice withdrawn from chronic cocaine. Physiol Behav 240, 113542. doi: 10.1016/J.PHYSBEH.2021.113542

44. Mañas-Padilla, M. C., Tezanos, P., Cintado, E., Vicente, L., Sánchez-Salido, L., Gil-Rodríguez, S., et al. (2023). Environmental enrichment alleviates cognitive and psychomotor alterations and increases adult hippocampal neurogenesis in cocaine withdrawn mice. Addiction Biology 28, e13244. doi: 10.1111/ADB.13244

45. Margolis, K. G., Cryan, J. F., and Mayer, E. A. (2021). The Microbiota-Gut-Brain Axis: From Motility to Mood. Gastroenterology 160, 1486–1501. doi: 10.1053/J.GASTRO.2020.10.066

46. Matsuoka, Y., Nakasone, H., Kasahara, R., and Fukuchi, M. (2024). Expression Profiles of Brain-Derived Neurotrophic Factor Splice Variants in the Hippocampus of Alzheimer’s Disease Model Mouse. Biol Pharm Bull 47, 1858–1867. doi: 10.1248/BPB.B24-00446

47. Matthews, K., and Robbins, T. W. (2003). Early experience as a determinant of adult behavioural responses to reward: the effects of repeated maternal separation in the rat. Neurosci Biobehav Rev 27, 45–55. doi: 10.1016/s0149-7634(03)00008-3

48. McWhirt, J., Sathyanesan, M., Sampath, D., and Newton, S. S. (2019). Effects of restraint stress on the regulation of hippocampal glutamate receptor and inflammation genes in female C57BL/6 and BALB/c mice. Neurobiol Stress 10, 100169. doi: 10.1016/J.YNSTR.2019.100169

49. Melgar-Locatelli, S., Mañas-Padilla, M. C., Castro-Zavala, A., Rivera, P., del Carmen Razola-Díaz, M., Monje, F. J., et al. (2024a). Diet enriched with high-phenolic cocoa potentiates hippocampal brain-derived neurotrophic factor expression and neurogenesis in healthy adult micewith subtle effects on memory. Food Funct 15, 8310–8329. doi: 10.1039/D4FO01201A

50. Melgar-Locatelli, S., Mañas-Padilla, M. C., Gavito, A. L., Rivera, P., Rodríguez-Pérez, C., Castilla-Ortega, E., et al. (2024b). Sex-specific variations in spatial reference memory acquisition: Insights from a comprehensive behavioral test battery in C57BL/6JRj mice. Behavioural brain research 459. doi: 10.1016/J.BBR.2023.114806

51. Millstein, R. A., and Holmes, A. (2007). Effects of repeated maternal separation on anxiety- and depression-related phenotypes in different mouse strains. Neurosci Biobehav Rev 31, 3–17. doi: 10.1016/J.NEUBIOREV.2006.05.003

52. Mitrea, L., Nemeş, S. A., Szabo, K., Teleky, B. E., and Vodnar, D. C. (2022). Guts Imbalance Imbalances the Brain: A Review of Gut Microbiota Association With Neurological and Psychiatric Disorders. Front Med (Lausanne) 9, 706. doi: 10.3389/FMED.2022.813204/BIBTEX

53. Nikolova, V. L., Hall, M. R. B., Hall, L. J., Cleare, A. J., Stone, J. M., and Young, A. H. (2021). Perturbations in Gut Microbiota Composition in Psychiatric Disorders: A Review and Meta-analysis. JAMA Psychiatry 78, 1343–1354. doi: 10.1001/JAMAPSYCHIATRY.2021.2573

54. Paxinos, G., and Franklin, K. B. L. (2012). The mouse brain in stereotaxic coordinates., 4th Edn. San Diego: Academic Press.

55. Peirce, J. M., and Alviña, K. (2019). The role of inflammation and the gut microbiome in depression and anxiety. J Neurosci Res 97, 1223–1241. doi: 10.1002/jnr.24476

56. Planchez, B., Surget, A., and Belzung, C. (2019). Animal models of major depression: drawbacks and challenges. J Neural Transm (Vienna) 126, 1383–1408. doi: 10.1007/s00702-019-02084-y

57. Portero-Tresserra, M., Gracia-Rubio, I., Cantacorps, L., Pozo, O. J., Gómez-Gómez, A., Pastor, A., et al. (2018). Maternal separation increases alcohol-drinking behaviour and reduces endocannabinoid levels in the mouse striatum and prefrontal cortex. European Neuropsychopharmacology 28, 499–512. doi: 10.1016/j.euroneuro.2018.02.003

58. Prospero, L., Riezzo, G., Linsalata, M., Orlando, A., D’Attoma, B., Di Masi, M., et al. (2021). Somatization in patients with predominant diarrhoea irritable bowel syndrome: the role of the intestinal barrier function and integrity. BMC Gastroenterol 21. doi: 10.1186/S12876-021-01820-7

59. Ramos, A., Pereira, E., Martins, G. C., Wehrmeister, T. D., and Izídio, G. S. (2008). Integrating the open field, elevated plus maze and light/dark box to assess different types of emotional behaviors in one single trial. Behavioural Brain Research 193, 277–288. doi: 10.1016/J.BBR.2008.06.007

60. Récamier-Carballo, S., Estrada-Camarena, E., and López-Rubalcava, C. (2017). Maternal separation induces long-term effects on monoamines and brain-derived neurotrophic factor levels on the frontal cortex, amygdala, and hippocampus: differential effects after a stress challenge. Behavioural pharmacology 28, 545–557. doi: 10.1097/FBP.0000000000000324

61. Ronquillo, J., Nguyen, M. T., Rothi, L. Y., Bui-Tu, T. D., Yang, J., and Halladay, L. R. (2023). Nature and nurture: Comparing mouse behavior in classic versus revised anxiety-like and social behavioral assays in genetically or environmentally defined groups. Genes Brain Behav 22, e12869. doi: 10.1111/GBB.12869

62. Rostami-Faradonbeh, N., Amini-Khoei, H., Zarean, E., Bijad, E., and Lorigooini, Z. (2024). Anethole as a promising antidepressant for maternal separation stress in mice by modulating oxidative stress and nitrite imbalance. Scientific Reports 2024 14:1 14, 1–10. doi: 10.1038/s41598-024-57959-2

63. Salta, E., Lazarov, O., Fitzsimons, C. P., Tanzi, R., Lucassen, P. J., and Choi, S. H. (2023). Adult hippocampal neurogenesis in Alzheimer’s disease: A roadmap to clinical relevance. Cell Stem Cell 30, 120–136. doi: 10.1016/J.STEM.2023.01.002

64. San Felipe, D., Martín-Sánchez, B., Zekri-Nechar, K., Moya, M., Llorente, R., Zamorano-León, J. J., et al. (2024). Consequences of Early Maternal Deprivation on Neuroinflammation and Mitochondrial Dynamics in the Central Nervous System of Male and Female Rats. Biology (Basel) 13, 1011. doi: 10.3390/BIOLOGY13121011/S1

65. Socała, K., Doboszewska, U., Szopa, A., Serefko, A., Włodarczyk, M., Zielińska, A., et al. (2021). The role of microbiota-gut-brain axis in neuropsychiatric and neurological disorders. Pharmacol Res 172, 105840. doi: 10.1016/J.PHRS.2021.105840

66. Sochocka, M., Diniz, B. S., and Leszek, J. (2016). Inflammatory Response in the CNS: Friend or Foe? Molecular Neurobiology 2016 54:10 54, 8071–8089. doi: 10.1007/S12035-016-0297-1

67. Sohrabji, F., and Lewis, D. K. (2006). Estrogen-BDNF interactions: implications for neurodegenerative diseases. Front Neuroendocrinol 27, 404–414. doi: 10.1016/J.YFRNE.2006.09.003

68. Stevens, B. R., Goel, R., Seungbum, K., Richards, E. M., Holbert, R. C., Pepine, C. J., et al. (2018). Increased human intestinal barrier permeability plasma biomarkers zonulin and FABP2 correlated with plasma LPS and altered gut microbiome in anxiety or depression. Gut 67, 1555. doi: 10.1136/GUTJNL-2017-314759

69. Sung, P. S., Lin, P. Y., Liu, C. H., Su, H. C., and Tsai, K. J. (2020). Neuroinflammation and Neurogenesis in Alzheimer’s Disease and Potential Therapeutic Approaches. Int J Mol Sci 21, 701. doi: 10.3390/IJMS21030701

70. Tao, E., Wu, Y., Hu, C., Zhu, Z., Ye, D., Long, G., et al. (2023). Early life stress induces irritable bowel syndrome from childhood to adulthood in mice. Front Microbiol 14. doi: 10.3389/FMICB.2023.1255525/FULL

71. Thevaranjan, N., Puchta, A., Schulz, C., Naidoo, A., Szamosi, J. C., Verschoor, C. P., et al. (2017). Age-Associated Microbial Dysbiosis Promotes Intestinal Permeability, Systemic Inflammation, and Macrophage Dysfunction. Cell Host Microbe 21, 455–466.e4. doi: 10.1016/J.CHOM.2017.03.002

72. Valvassori, S. S., Varela, R. B., and Quevedo, J. (2017). “Animal Models of Mood Disorders: Focus on Bipolar Disorder and Depression,” in Animal Models for the Study of Human Disease: Second Edition, (Elsevier Inc.), 991–1001. doi: 10.1016/B978-0-12-809468-6.00038-3

73. Varanoske, A. N., McClung, H. L., Sepowitz, J. J., Halagarda, C. J., Farina, E. K., Berryman, C. E., et al. (2022). Stress and the gut-brain axis: Cognitive performance, mood state, and biomarkers of blood-brain barrier and intestinal permeability following severe physical and psychological stress. Brain Behav Immun 101, 383–393. doi: 10.1016/J.BBI.2022.02.002

74. Veenit, V., Zhang, X., Ambrosini, A., Sousa, V., and Svenningsson, P. (2021). The Effect of Early Life Stress on Emotional Behaviors in GPR37KO Mice. Int J Mol Sci 23. doi: 10.3390/IJMS23010410

75. Wei, R. M., Zhang, Y. M., Feng, Y. Z., Zhang, K. X., Zhang, J. Y., Chen, J., et al. (2023). Resveratrol ameliorates maternal separation-induced anxiety- and depression-like behaviors and reduces Sirt1-NF-kB signaling-mediated neuroinflammation. Front Behav Neurosci 17. doi: 10.3389/FNBEH.2023.1172091/FULL

76. Yu, H., Lin, L., Zhang, Z., Zhang, H., and Hu, H. (2020). Targeting NF-κB pathway for the therapy of diseases: mechanism and clinical study. Signal Transduction and Targeted Therapy 2020 5:1 5, 1–23. doi: 10.1038/s41392-020-00312-6

77. Zaghloul, N., Kurepa, D., Bader, M. Y., Nagy, N., and Ahmed, M. N. (2020). Prophylactic inhibition of NF-κB expression in microglia leads to attenuation of hypoxic ischemic injury of the immature brain. J Neuroinflammation 17, 1–13. doi: 10.1186/S12974-020-02031-9/FIGURES/8

